# Using graph-based model to identify cell specific synthetic lethal effects

**DOI:** 10.1101/2023.07.23.550246

**Authors:** Mengchen Pu, Kaiyang Cheng, Xiaorong Li, Yucui Xin, Lanying Wei, Sutong Jin, Weisheng Zheng, Gongxin Peng, Qihong Tang, Jielong Zhou, Yingsheng Zhang

## Abstract

Synthetic lethal (SL) pairs are pairs of genes whose simultaneous loss-of-function results in cell death, while a damaging mutation of either gene alone does not affect the cell’s survival. This makes SL pairs attractive targets for precision cancer therapies, as targeting the unimpaired gene of the SL pair can selectively kill cancer cells that already harbor the impaired gene. Limited by the difficulty of finding true SL pairs, especially on specific cell types, the identification of SL targets still relies on expensive, time-consuming experimental approaches. In this work, we utilized various cell-line specific omics data to design a deep learning model for predicting SL pairs on particular cell-lines. By incorporating multiple types of cell-specific omics data with a self-attention module, we represent gene relationships as graphs. Our approach demonstrates the potential to facilitate the discovery of cell-specific SL targets for cancer therapeutics, providing a tool to unearth mechanisms underlying the origin of SL in cancer biology. Our approach allows for prediction of SL pairs in a cell-specific manner and enhances cancer precision medicine. The code and data of our approach can be found at https://github.com/promethiume/SLwise

**Highlights:**

- Few computational methods can systematically predict SL pairs at a cell-specific level, and their performance may not generalize well to clinical scenarios due to the heterogeneity of cancer types.
- The SLWise utilizes various cell-line specific omics data to design a deep learning model with a graph-based representation and self-attention mechanism.
- This approach allows for the prediction of SL pairs in a cell-specific manner, providing valuable insights on effectively identifying the cell-type specific SL targets for personalized treatment strategies.

## 1. Introduction

Over the past decade, precision medicine has gained widespread acceptance as a concept for developing targeted therapies based on individual biological background. Identification of molecular biomarkers is now a common practice in clinical studies, especially in the field of cancer therapy. Synthetic lethality (SL), where simultaneous inactivation of a gene pair causes cell death, is considered to be of significant importance in cancer treatment. Cancer cells [1] often have a large number of damaging mutations and gene replication errors that are not present in normal cells. If the corresponding SL gene pair of a cancer cell is found as a target, it is possible to precisely kill tumors with the specific mutation without damaging healthy cells. The SL mechanism [2] has the potential to be utilized in precision, anti-cancer drug development. The availability of high-throughput genomics data and therapeutic agents makes cancer an ideal field for the study of precision medicine, which matches a patient’s biological background with the selection of target-oriented drugs. However, identifying SL pairs with large scaled *in vitro* experimental screening is time- and labor-consuming. Thus, an accurate *in silico* predictor for SL pairs is deemed necessary [3].

Considering the limitations of experiments and the complexity of the SL mechanism, researchers have undertaken efforts to develop computational models for predicting SL in a more efficient manner. Machine learning (ML) and deep learning (DL) methods can effectively integrate multi-dimensional biological data such as paralog data [4], mutation patterns [4, 5], expression profiles [6] and protein-protein interaction networks (PPI) [7]] etc., and then perform feature learning through parameter fitting. These methods distill decisive correlations from these comprehensive information data for reliable SL prediction. The GRSMF model [8] uses known SL interactions to learn the association representation and functional similarity from Gene Ontology (GO) to predict potential SL interactions in a pan-cancer cell manner. This is done by applying the information provided by known SL interactions and gene functional annotations. The SL2MF model [9] uses the SL interactions with additional GO similarity matrix and PPI similarity matrix as supporting information. It leverages the similarity in the network representation when two genes share similar functions. All the predicted SL pairs were ranked and compared with known SL pairs to evaluate the model’s prediction ability. Due to the large dimensionality of the SL gene interaction matrix, it is difficult for machine learning methods to capture the underlying relationship among them. In recent years, graph neural networks (GNNs) have been proven superior in the issue of link prediction. In the KG4SL model [10], the knowledge graph is used to introduce additional information related to genes, such as compounds, diseases, and biological processes, which improves the prediction performance. It is the first model that integrates a knowledge graph and a GNN for SL prediction. The NF4SL model [11] includes a strategy for enhancing gene representation by incorporating both global and local information. The model uses two graph-level modules for data enhancement and similarity learning between interacting genes to increase similarity between their representations. The GCATSL model [7] introduced the Graph Attention Network (GAT) [12] model for the first time to predict SL pairs, and also used additional data such as PPI and GO to construct separate feature maps as auxiliary information for the model. The GAT captures the local and global features of each gene node, using additional feature data to weigh and sum specific feature representations to ultimately reconstruct the predicted probability matrix. The MGE4SL model [13] uses extra data from sources such as Corum, Reactome, KEGG, and STRING. They employ a Graph Convolutional Networks (GCN) [14] encoder to obtain feature representations of the gene nodes and their neighboring nodes. It combines the features of SL pairs and additional knowledge graph features to obtain a mixed matrix representation of all gene information. The SLMGAE model [15] treated SL pairs as the main graph and the other data sources (e.g., PPI, GO, etc.) as the support view. The implementation of the self-attention mechanism by assigning a randomly initialized and normalized weight matrix to each view, which makes each view adaptively learn the relationship between features. Due to the uncertainty of the weight matrix, the association between features becomes scattered when more and more features are introduced, making it challenging to incorporate additional biological characteristics for genes or cells.

The context of species, tissue types, cell types and cellular conditions determines the SL interaction. This complex phenomenon known as the context-specific or context-dependent of SL pairs. Theoretically, by co-inactivating a cancer specific SL pair, a normal tissue can maintain its fitness and resist malignancy, as the cancer cells are selectively killed by the specific lethal effect. This might be dependent on certain intrinsic conditions such as the heterogeneity of different cell types, hypoxia, external disturbances which can result in specific genetic interaction networks [16, 17]. Thus, the synthetic lethal effects may vary depending on the tumor cells or cell lines. Besides, targeting tumor-specific or cell-line specific SL pairs could also help overcome the resistance to synthetic lethal drugs that target heterogeneous tumors as a whole, and has the potential for treating tumors with various complex conditions in future. Only few computational methods exist that are able to predict cell-line specific SL pairs. EXP2SL [5] is a semi-supervised cell-specific SL pairs predictor utilizing L1000 gene expression profiles. Each gene is represented by a 978-dimensional z-score of the shRNA perturbation profile. The extracted features using MLP layers for given gene pairs are concatenated to predict the SL confidence score. In the inference phase, genes with perturbation profile data are the only ones that can be used to make predictions. MVGCN-iSL [5, 18, 19] applies multi-view graph convolutional network (GCN) model to predict cell-specific SL pairs. Several cell-independent networks data, cell-dependent gene data and SL pairs information are utilized. The cell-specific relationship between genes is only provided by cell-specific SL labels, which has shown to be the most informative.

Despite having high prediction scores when using data from SL pairs database, the performance of state-of-the-art (SOTA) models may not generalize well on cell-line specific SL pairs. It remains challenging to identify new, robust cell-line specific SL pairs. Several studies rely on perturbation data [5, 18, 19], which is experimentally costly given the enormous number of gene sets and cell lines involved. It is difficult to generalize the results of perturbation experiments to all genes under all cell lines. Additionally, minor variations in tumor cell mutation profiles may exist between different clinical patients, it is deemed impractical to train a model for every single cell line in clinical practice. In this study, we have developed a computational tool to predict cell-specific SL pairs using cell-line specific omics data as input. Our approach employs a graph-based representation technique to represent the inter-relationship between genes comprehensively. Additionally, we incorporate a self-attention mechanism, which allows our model to identify the most relevant parts of the input data. We demonstrate that our predictor has the potential to be applied to diverse cancer cell-lines. Furthermore, we also evaluate our model on a cell-line transferable study, which determines how well the model can generalize to new cell-lines, and how accurately it can predict in unknown cell-lines.

## 2. Materials and method

### 2.1. Data pre-processing

To predict cell-line specific SL pairs, we applied individual cell-line multi-omics data as inputs. We used L1000 data, gene effect scores (ES), the exclusive mutation (EM) patterns in our approach. Besides, the 36600 paralog pairs were obtained directly from the supplementary data of Kegel et.al. [20].

The level 5 data of L1000 as the original analysis dataset was directly downloaded from the LINCS L1000 project [19]. We transformed the data into a matrix of 12328 rows and 238351 columns, where each row represents a gene, and each column represents the normalized expression fold change values of that gene after perturbation, compared to the control group, under various cell-lines. Based on this matrix and annotation information of the level 5 data, we used data from the CRISPR gene knock-out [21], setting the absolute value of the fold change threshold as 1.5. The columns in this matrix represents the cell-lines, the target genes, the perturb genes, and the fold change values.

Gene ES and gene expression profiles were obtained from the DepMap portal, the public 22Q2 dataset [22]. The gene ES were originally derived from CRISPR knockout screens conducted by Broad’s Project Achilles and Sanger’s SCORE projects [23-25], which reflect the normalized impact of knocking out a specific gene on a certain cell-line. Negative scores indicate cell growth inhibition and/or death following gene knockout, scores lower than -0.5 indicating depletion on most cell-lines. For each cell-line, we computed z-scores for both gene expression data and gene ES using all genes. We then identified genes with low expression or low ES for each cell-line, defined as genes with expression values < 1 and expression z-scores < -1.28 (corresponding to the lowest 10% of the standard normal distribution), or with ES lower than -0.5 and gene effect z-scores lower than -1.28.

The somatic mutations of the corresponding TCGA cohorts from cBioPortal database (https://www.cbioportal.org/) were utilized to generate mutually exclusive mutation patterns for all tested gene pairs in each cancer cell-line. We employed a weighted sampling-based approach, called WeSME [26], to identify significant mutually exclusive gene pairs. After creating a binary gene-sample mutation matrix for the same tumor cohort by recording whether a sample had one or more non-synonymous mutations in a certain gene, we calculated the mutation frequencies of samples, and estimating a null distribution of the mutation profiles of a gene by conducting a simulation 1000 times based on the mutation frequencies of samples. We deemed gene pairs with p-values less than 0.05 to be mutually exclusive.

GEMINI, a variational Bayesian method [27], was applied to generate the ground truth for our model. It was utilized to identify lethal interactions from high-throughput CRISPR-based combinatorial perturbation experiment results [28-31]. The sensitive interaction score was generated from GEMINI for each gene pair and the false discovery rate (FDR), sensitivity scores, and p-values were applied with 0.05 as cutoff for the final ground truth of positive SL pairs. Those pairs with negative SL scores and among the bottom 50% were used to define the negative ground truth. To address the imbalance between the positive and negative SL pairs for each cell-lines, we randomly selected a subset of the majority class with an equal number of examples as the minority class to ensure both positive and negative pairs were equally sampled during training and model evaluation. We employed FDR, p-values, and sensitive scores to identify positive labels from GEMINI calculation and integrated all the results as positive samples.

Finally, we obtained 173, 130, and 1345 positive samples and 1446, 28, and 2820 negative samples in HT29, A375, and A549 respectively. We balanced the negative and positive samples and partitioned the dataset into a training set (80%) and a validation set (20%), and then used four groups for training and one group for testing. The metric for evaluating multi-omics data and model architectures is the average performance of all the folds. In addition, we used SL pairs identified in recent years (from 2018 to 2021) in the same cell-line as an external test set to validate the model’s ability to make specific cell-line predictions.

### 2.2. Graph-based neural network

#### 2.2.1 Overview

The prediction task of SL interactions can be represented as a matrix completion task, which aims to predict unobserved interactions. The interactions between and within the different omics levels, such as genomics, transcriptomics, proteomics, and metabolomics, form vast and complex networks, which are challenging to understand. The graph neural network, on the other hand, is well-suited to handle such type of non-Euclidean data. This makes it an ideal tool for analyzing and understanding the complex network of interactions within the omics levels. Here, we presented a graph-based model for SL prediction. The framework of our approach is illustrated in Figure 1. It uses graphs generated from multiple biological data of gene and cell information, with the SL graph serving as the reference in training process. To generate relevant features, we employ various graph encoders to extract features that take different perspectives on the data, and utilize a self-attention mechanism to integrate all the reconstructed graphs. This is then followed by using a multi-layer perceptron (MLP) for SL pair prediction. In following sections, each module in our approach is detailed. The symbols and notations used are summarized in Table 1.

**Table 1.**
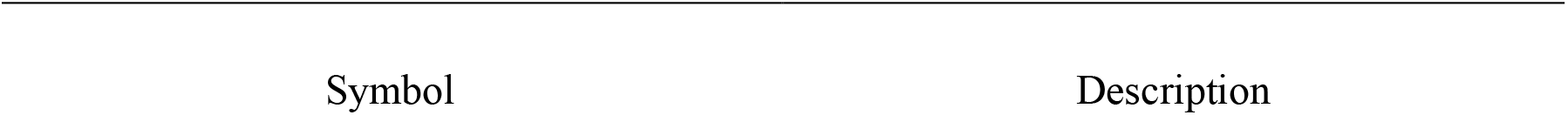
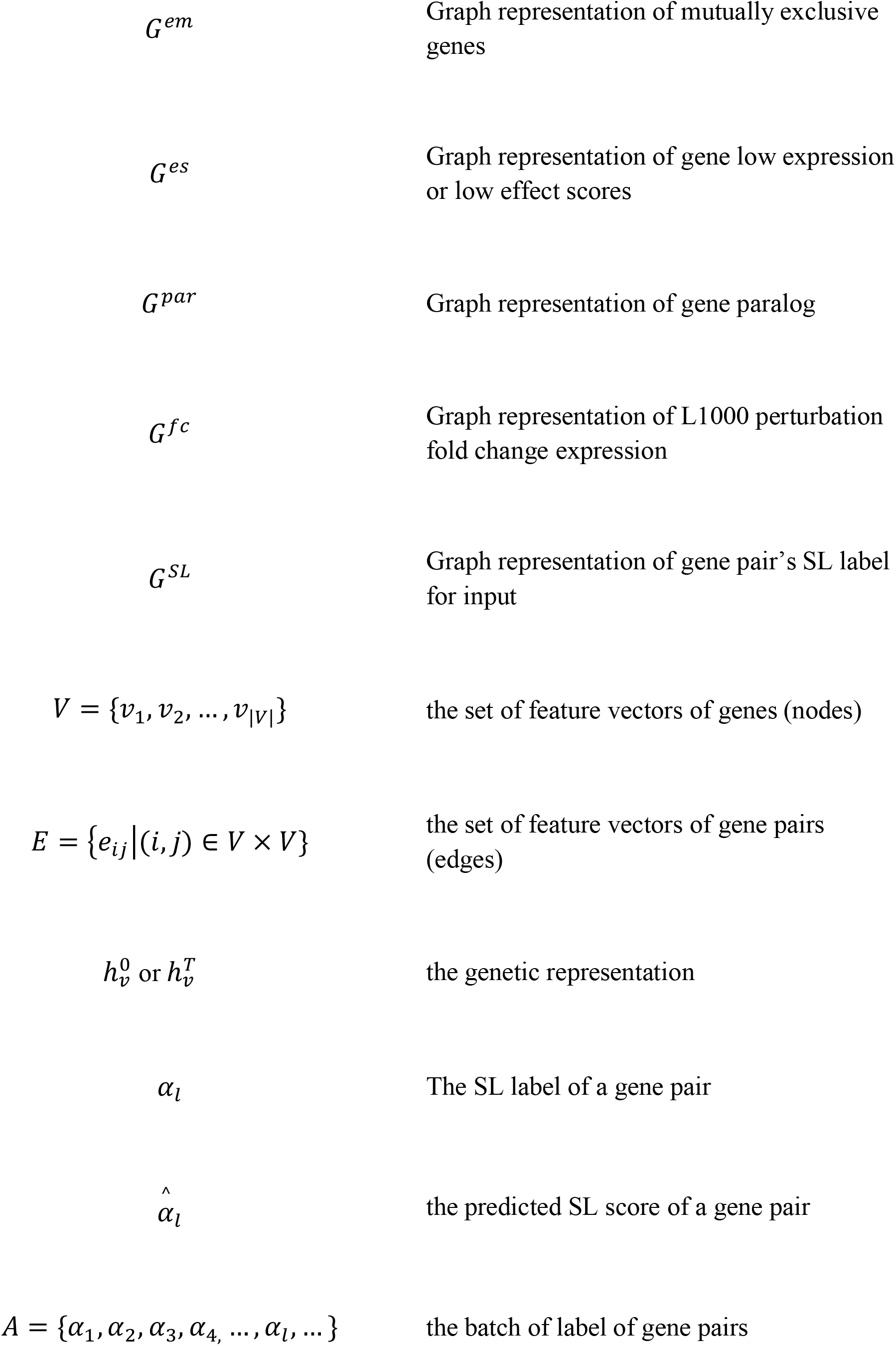
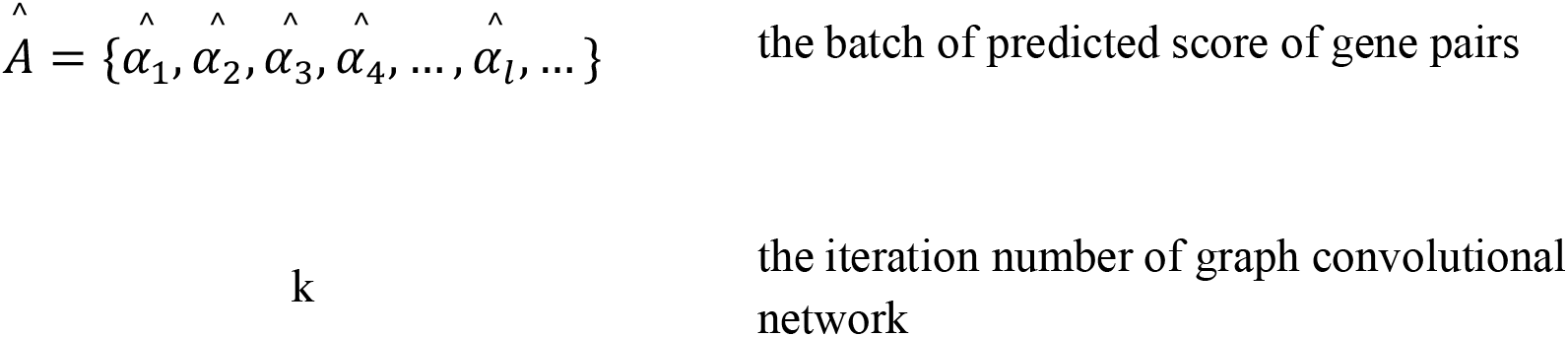
List of symbols and notations used in the paper.

**Figure 1.**
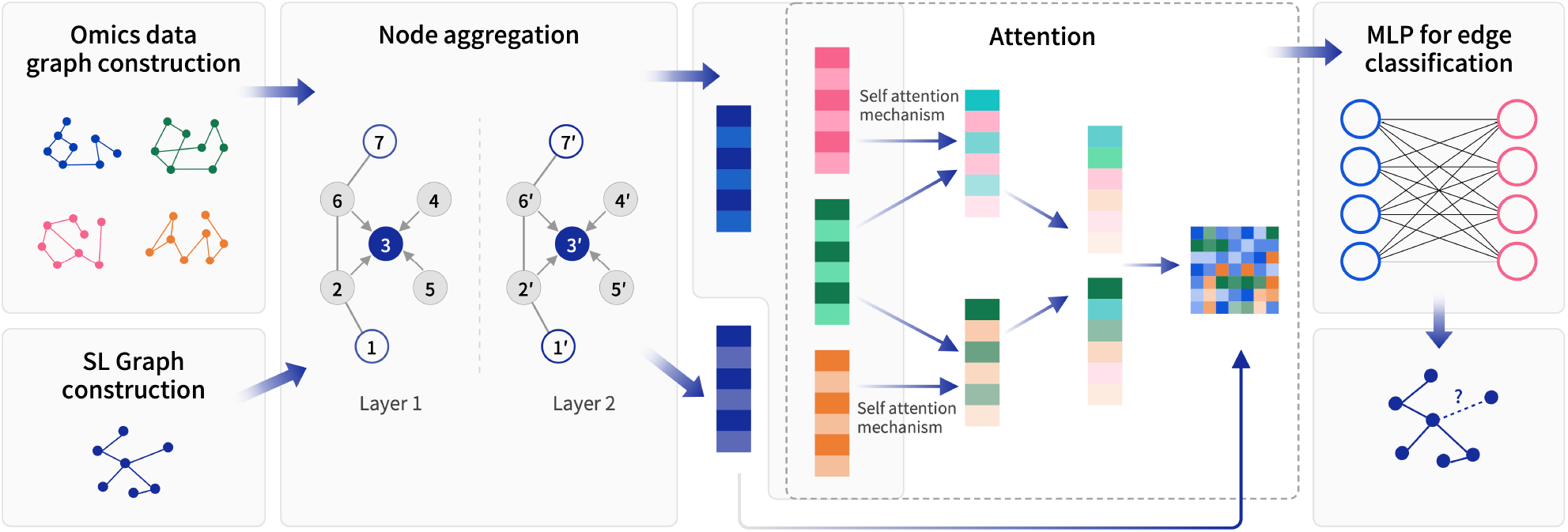
The framework of our method.

#### 2.2.2 Graph neural network for cell specific feature extraction

To better understand the relationship between genes in distinguished cells, we applied multi-omics data that are converted to a graph representation *G*^*em*^, *G*^*es*^, *G*^*par*^, and *G*^*fc*^respectively. We first convert the omics data into a graph representation, in which *G*={*v*_),_ *v*_*,_…,*v*_|,|_} is the set of the set of feature vectors of genes, *E*={*e*_*ij*_|(*i,j*{ ∈ *V* × *V*} is the set of feature vectors of a pair of genes, and *e*_*ij*_ ∈ {0,1}^*m*^ where *m* is the number of cell-line genes and 1 indicates the whether the value of *gene*_*i*_ and *gene*_*j*_ in those data matrices is 1.

We built a two-layer GraphSAGE [32] module with a fixed number of sampled neighbors to aggregate information for each omics data, since two-hops neighborhood covers most connection information and is usually sufficient for large graph structure learning. The different dimensions of two graph convolution modules are 128 and 16, respectively, each operation is defined as:

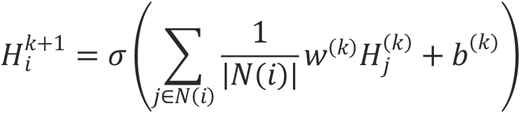

Where *N*(*i*) is the neighbor gene list, *w*^(*k*)^ is the trainable parameter matrix of the k-th layer, 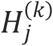 is the representation matrix of gene *j, b*^(*k*)^ is the bias term at layer *k*, and σ is the activation function. The rectified linear unit (ReLU) is applied, which is defined as follows:

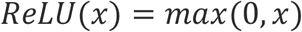

To avoid overfitting, we added a dropout function after each convolutional block with a probability of 0.5.

The complexity and homogeneity of those omics data make it crucial to assign specific edge weights in order to accurately identify important genes. In this case, we assigned a weight of 1 to the EM data, the ES data and paralog data, due to their binary nature. For the L1000 data, which represent gene expression in treatment, we used the actual numbers from the data matrix as edge weights. To prevent over-smoothing in the graph neural network, we only considered edge weights with values greater than three or less than negative three, effectively eliminating less important nodes.

To generate a more informative embedding, we applied an aggregation operation, *z*_*s*_, to ensemble cell specific representation of genes representation in multi-omics derived from different graph structures. the final embedding can be represented as

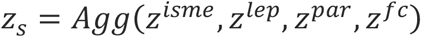

Here, we simply concatenate these latent features together for use in the next attention module.

#### 2.2.3. Feature fusion module

We present a multi-head transformer cross-attention method that directs attention to three features such as EM data from both ES data and paralogs in two stages. The three omics data (EM, ES, paralog) are derived from encoders and then fed into the multi-head attention module [33], also known as the transformer block. The latent feature, combined with SL pairs feature representation, is passed through an MLP layer for SL interactions reconstruction. We use the attention mechanism to learn the weight distribution of different features, which helps to identify the important features for prediction. The multi-head attention is calculated by the following formulas

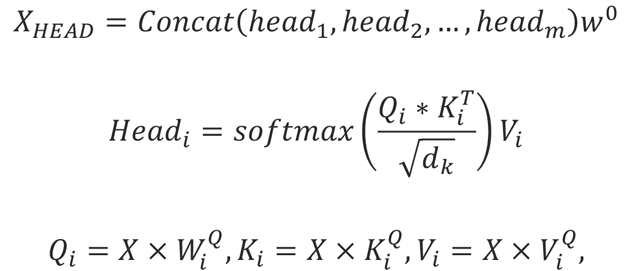

where *Q*_*i*_, *K*_*i*_ and *V*_*i*_ are the Q, K and V matrices derived from the linear transformation of those biological features are passed through the attention layer and the feed-forward network layer. For the L1000 data, we simply concatenated the GraphSAGE features of it with the output of the attention module.

The final representation for link prediction (positive SL pairs) is created by combining that relevant information with the graph representations of SL pairs, *GSL*. Additionally, layer normalization is also applied to accelerate the convergence of the neural network and prevent the ‘covariate-shift’ and ‘high-parameters’ issues.

#### 2.2.4. MLP for edge prediction

We implemented a fully connected neuron network to predict the potential SL pairs. It takes in features from the fusion layer as input and has multiple hidden layers utilizing ReLU as activation function. The output layer contains a single node with a sigmoid activation function, which outputs a probability indicating the likelihood that a given edge corresponds to an SL pair in a specific cell-line. The Binary Cross Entropy loss function is applied.

The training process of our approach is illustrated in Algorithm 1.

#### 2.2.5 Ablation experiment

In our experiments, we evaluated the impact of specific combinations of input features and architectural components on our current model by conducting feature ablation and module ablation. In these experiments, we removed individual input features from multi-omics data in feature ablation and replaced different feature extraction modules or feature aggregation modules in module ablation.

Besides GraphSAGE, other graph based encoders such as GCN, GAT [12], and Topology adaptive graph convolutional networks (TAG) [34] are also designed for processing graph-structured data through the use of aggregation function, combination function, and readout function. GraphSAGE learns node features by aggregating information from a fixed number of sampled neighbors. GCN [14] use convolutional operations to aggregate information from neighboring nodes, GATs use attention mechanisms to weight the importance of the neighboring nodes while aggregating the node’s feature information. while TAG is a variant of GCN that adapts its convolutional filter based on the graph’s topology. These three architectures are utilized in the ablation study of feature extraction module.

The final model incorporated an attention mechanism into the omics feature aggregation modules. Due to the time and computational complexity of the transformer module, we conducted an additional ablation experiment in which we substituted the transformer module with a much simpler operation that concatenated the output of GraphSAGE modules. This experiment demonstrated the impact of the transformer module on the model performance.

#### 2.2.6 Performance evaluation

To evaluate the performance of our approach, we use four metrics: recall, precision, the area under the precision-recall curve (AUPR) and the area under the receiver operating characteristic curve (AUROC). We compare our predictor with EXP2SL, which is a cell-line specific predictor that serves as a baseline. All the metrics were calculated using Python (Scikit Learn package). The sparsity of the SL labels may lead to overfitting of the model. To address this issue, we conducted three types of evaluations with different ways of splitting the dataset. In evaluation study 1 (CV1), the dataset was partitioned by gene pairs, such that both genes in a test set might also appear in the training set. In evaluation study 2 (CV2), we divided the dataset by genes, ensuring that only one gene in a test pair was also present in the training set. In evaluation study 3 (CV3), we separated the dataset by genes, excluding both genes in a test set from the training set.

## 3. Results And Discussion

### 3.1. Overall performance

We evaluated our approach using five-fold cross-validation with specific settings on the aforementioned datasets for the three cell-lines and then compared the performance of our model to the SOTA EXP2SL model as baseline. We started by splicing the ground truth data into training and testing sets, and assessing the model’s performance within a single cell-line using AUC, AUPR, recall and precision, which presented in Table 2. Compared to the baselines, our model demonstrated a relatively stable performance with only a slight decline from CV1 to CV3 test, whereas the performance of the EXP2SL model declined significantly.

**Table 2.**
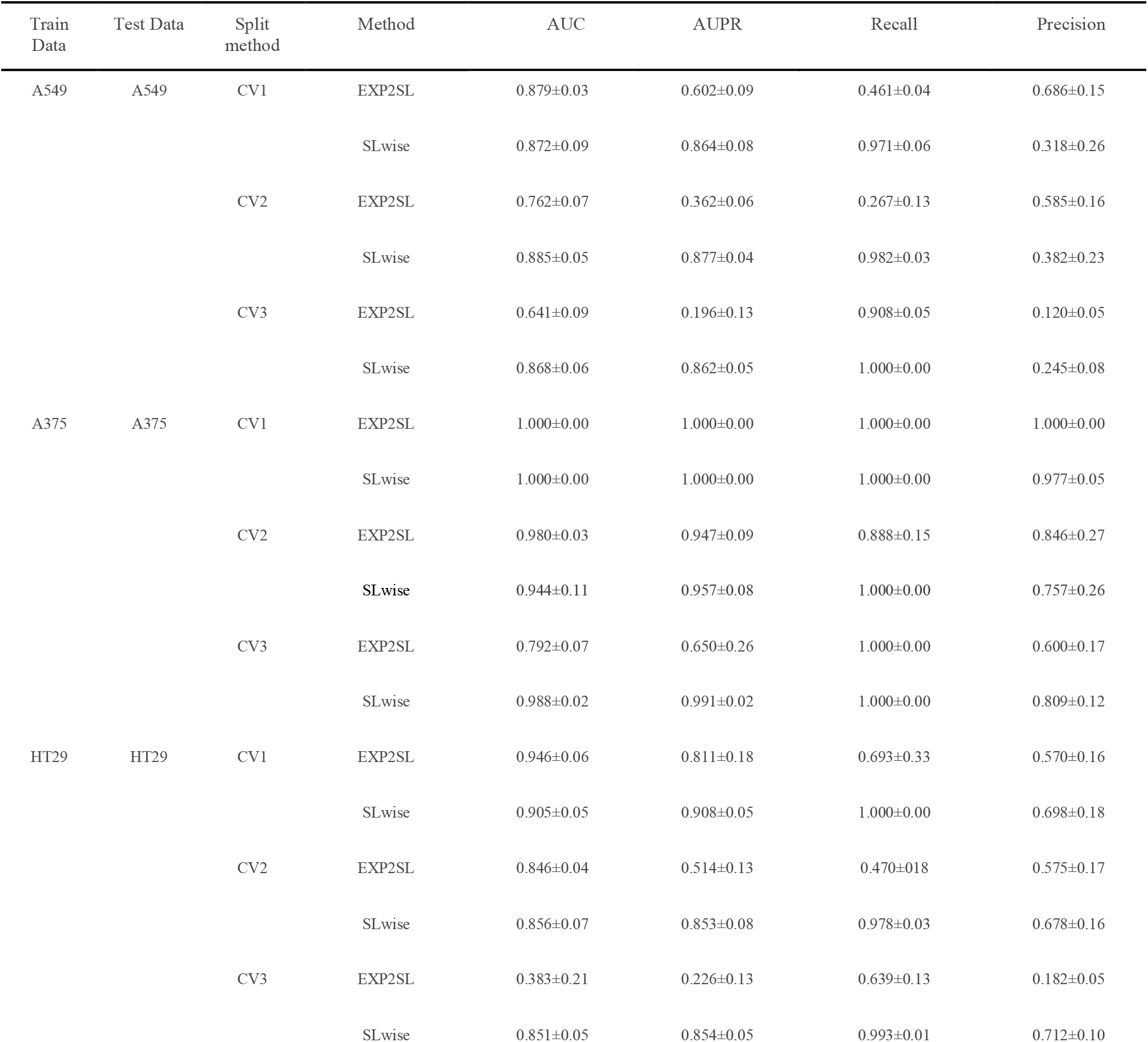
The performance of evaluation in three different cell lines under different split test set.

| Train Data | Test Data | Split method | Method | AUC | AUPR | Recall | Precision |
| --- | --- | --- | --- | --- | --- | --- | --- |
| A549 | A549 | CV1 | EXP2SL | 0.879±0.03 | 0.602±0.09 | 0.461±0.04 | 0.686±0.15 |
|  |  |  | SLwise | 0.872±0.09 | 0.864±0.08 | 0.971±0.06 | 0.318±0.26 |
|  |  | CV2 | EXP2SL | 0.762±0.07 | 0.362±0.06 | 0.267±0.13 | 0.585±0.16 |
|  |  |  | SLwise | 0.885±0.05 | 0.877±0.04 | 0.982±0.03 | 0.382±0.23 |
|  |  | CV3 | EXP2SL | 0.641±0.09 | 0.196±0.13 | 0.908±0.05 | 0.120±0.05 |
|  |  |  | SLwise | 0.868±0.06 | 0.862±0.05 | 1.000±0.00 | 0.245±0.08 |
| A375 | A375 | CV1 | EXP2SL | 1.000±0.00 | 1.000±0.00 | 1.000±0.00 | 1.000±0.00 |
|  |  |  | SLwise | 1.000±0.00 | 1.000±0.00 | 1.000±0.00 | 0.977±0.05 |
|  |  | CV2 | EXP2SL | 0.980±0.03 | 0.947±0.09 | 0.888±0.15 | 0.846±0.27 |
|  |  |  | SLwise | 0.944±0.11 | 0.957±0.08 | 1.000±0.00 | 0.757±0.26 |
|  |  | CV3 | EXP2SL | 0.792±0.07 | 0.650±0.26 | 1.000±0.00 | 0.600±0.17 |
|  |  |  | SLwise | 0.988±0.02 | 0.991±0.02 | 1.000±0.00 | 0.809±0.12 |
| HT29 | HT29 | CV1 | EXP2SL | 0.946±0.06 | 0.811±0.18 | 0.693±0.33 | 0.570±0.16 |
|  |  |  | SLwise | 0.905±0.05 | 0.908±0.05 | 1.000±0.00 | 0.698±0.18 |
|  |  | CV2 | EXP2SL | 0.846±0.04 | 0.514±0.13 | 0.470±0.18 | 0.575±0.17 |
|  |  |  | SLwise | 0.856±0.07 | 0.853±0.08 | 0.978±0.03 | 0.678±0.16 |
|  |  | CV3 | EXP2SL | 0.383±0.21 | 0.226±0.13 | 0.639±0.13 | 0.182±0.05 |
|  |  |  | SLwise | 0.851±0.05 | 0.854±0.05 | 0.993±0.01 | 0.712±0.10 |

Table 2 summarizes the performance of our approach and EXP2SL on three different cell-lines: A375, A549, and HT29. Both methods achieved similar results under CV1, where they both performed best on A375 among other cell-lines. However, our approach showed a clear advantage over EXP2SL under CV2 and CV3, where the gene sets for training and testing are more independent. Under CV2 datasets, our model improved by 12.3% and 1% on AUC, 51.5% and 33.9% on AUPR compared to EXP2SL in A549 and HT29 respectively. In particular, under CV3, EXP2SL suffers from a significant decline in performance, while our approach maintained a high level across all three cell-lines. Our model improved by 51%, and 30.7% on Recall, and by 26.3%, and 9.7% on AUPR in the A549, and HT29 respectively.

This demonstrates that our model is able to effectively capture the cell features from multi omics data and predict SL pairs in the dataset for different cell lines. It can be noticed that our approach is also a little bit struggled in identifying SL pairs within the cell-line A549. However, the overall performance of this study suggests that this is a promising approach for identifying SL pairs in genetic datasets.

### 3.2. Ablation result

The ablation study helps to verify the rationality of our current features and model architecture, and provide valuable insights into how to improve the model in the future. In this paper, we investigate the impact of the encoder module, the attention module and input omics data on the accuracy of predicting SL interactions. The results are shown in Supplementary Table S1-S3.

We conducted an ablation study to evaluate the performance of different encoder modules in our model for predicting SL interactions. We compared three types of graph convolutional network encoders (GAT, GCN, and TAG) and validated our model in three cell lines, averaging the results and compared with our original selection, GraphSAGE. As shown in **Table S1**, the performance of the GraphSAGE encoder module was slightly inferior to GCN in terms of AUC and AUPR, but demonstrated a competitive advantage in other metrics, showing that mean aggregation from neighborhood features of each node from multi-omics graph is very effective for SL prediction. Moreover, the fixed-number neighborhood sampling strategies enable GraphSAGE to aggregate a subset of each gene node, making it more scalable.

In addition, we also conducted an ablation study to evaluate the effect of attention mechanism on the feature aggregation module. The experimental results (Table S2) demonstrate that incorporating attention module can significantly improve the model performance compared with simply concatenating the extracted features.

Furthermore, we performed ablation study to examine the impact of different multi-omics data on effectiveness of our model. Details of the performance on each cell-line are shown in Table S3. We tested different combinations of input data by removing one of those different omics data types, and compared the result with those of the complete dataset. The ablation results demonstrate that using the complete dataset, the performance shows improvements on all metrices, indicating that each multi-omics data is beneficial to predict known positive SL samples in our approach. These experiments show that utilizing all multi-omics information to predict SL interactions can improve the performance of predictions. In the future, we may incorporate additional omics data or use data enhancement techniques to improve our predictions even further.

### 3.3. Cell-line transferable study

Finally, we evaluated the model’s ability to predict SL pairs in unknown cell-lines, where the training and testing data came from different cell-lines, mimicking a real-world scenario. It helps to establish the model’s ability to be used in practical applications. The preliminary results appear to be promising when taking into account the number of SL label for each cell-lines.

Our model was able to make accurate predictions when using data from different cell-lines for training and testing (Table 3). Our model had the best performance when using A549 as training data. The large amount of single-label (SL) data available for A549, 1345 positives and 2820 negatives, may have contributed to its performance. This dataset is almost five times bigger than the amount available for the other two cell lines (A375 and HT29). Besides, our approach also managed to maintain a satisfactory level of performance when using the other cell-lines as training data. In these tests, EXP2SL performed poorly. These results suggest that our model is transferable among cell lines, and has the potential to predict SL pairs for unknown cell-lines.

**Table 3.** The performance of transferable evaluation in three different cell lines under different split test set.

| Train Data | Test Data | Method | AUC | AUPR | Recall | Precision |
| --- | --- | --- | --- | --- | --- | --- |
| A549 | A375 | EXP2SL | 0.852±0.02 | 0.773±0.04 | 0.400±0.08 | 0.853±0.11 |
|  |  | SLwise | 0.902±0.04 | 0.960±0.02 | 0.811±0.11 | 0.954±0.08 |
|  | HT29 | EXP2SL | 0.708±0.04 | 0.343±0.14 | 0.311±0.13 | 0.400±0.21 |
|  |  | SLwise | 0.962±0.02 | 0.986±0.01 | 0.847±0.10 | 0.938±0.09 |
| A375 | A549 | EXP2SL | 0.545±0.03 | 0.165±0.01 | 0.177±0.00 | 0.128±0.02 |
|  |  | SLwise | 0.822±0.01 | 0.672±0.01 | 0.728±0.10 | 0.562±0.17 |
|  | HT29 | EXP2SL | 0.641±0.04 | 0.276±0.1 | 0.589±0.13 | 0.180±0.03 |
|  |  | SLwise | 0.925±0.03 | 0.973±0.01 | 0.812±0.11 | 0.945±0.08 |
| HT29 | A549 | EXP2SL | 0.608±0.03 | 0.166±0.02 | 0.118±0.00 | 0.202±0.04 |
|  |  | SLwise | 0.822±0.00 | 0.713±0.01 | 0.610±0.08 | 0.744±0.01 |
|  | A375 | EXP2SL | 0.707±0.10 | 0.575±0.06 | 0.367±0.05 | 0.557±0.09 |
|  |  | SLwise | 0.953±0.01 | 0.981±0.01 | 0.765±0.01 | 1.00±0.00 |

## 4. Conclusion

We presented a novel method, SLWise, that combines graph-based representations, attention mechanisms, and multiple omics data to enhance its capabilities. The ablation study shows that the GraphSAGE module effectively captures the representation of omics data. The transformer cross-attention mechanism is designed to assemble multi-source features, making it better at capturing the cell specific correlation of data or features. By integrating different biological data sources, our model can capture the complex relationships and interactions within the data, and thus outperform SOTA models in predicting cell-specific SL pairs for different cell-lines. The development of our approach is expected to be beneficial to the advancement of cancer precision medicine by supporting the discovery of cell-type specific drug targets and biomarkers in the future.

## Supporting information

Supplemental Table1_SigProfiler_30-denovo-sigs

## Acknowledgements

We thank Dr. B. Fu and Dr. K. Tian from the Innovation Center for helpful discussions. We also thank Y. Wang, S. Guo, X. Xiang, and all members of the StoneWise AI. Inc. for administrative support.

## Author contributions statement

M.P. and Y.Z. conceived the project. M.P., K.C. and X.L. designed the method and conducted the experiments. W.Z, L.W. G.P. and Y.X. curated the data. M.P., K.C. and X.L. pre-processed the data and analysed the results. S. J. assisted with data analysis. Q.T. assisted with data visualizations. M.P., X.L. and K.C. wrote the manuscript. All authors provided feedback on the manuscript.

## Data Availability

The data underlying this article are available in the article and in its online supplementary material. The code and training data is available in https://github.com/promethiume/SLwise

### Algorithm 1. The training process of our model.

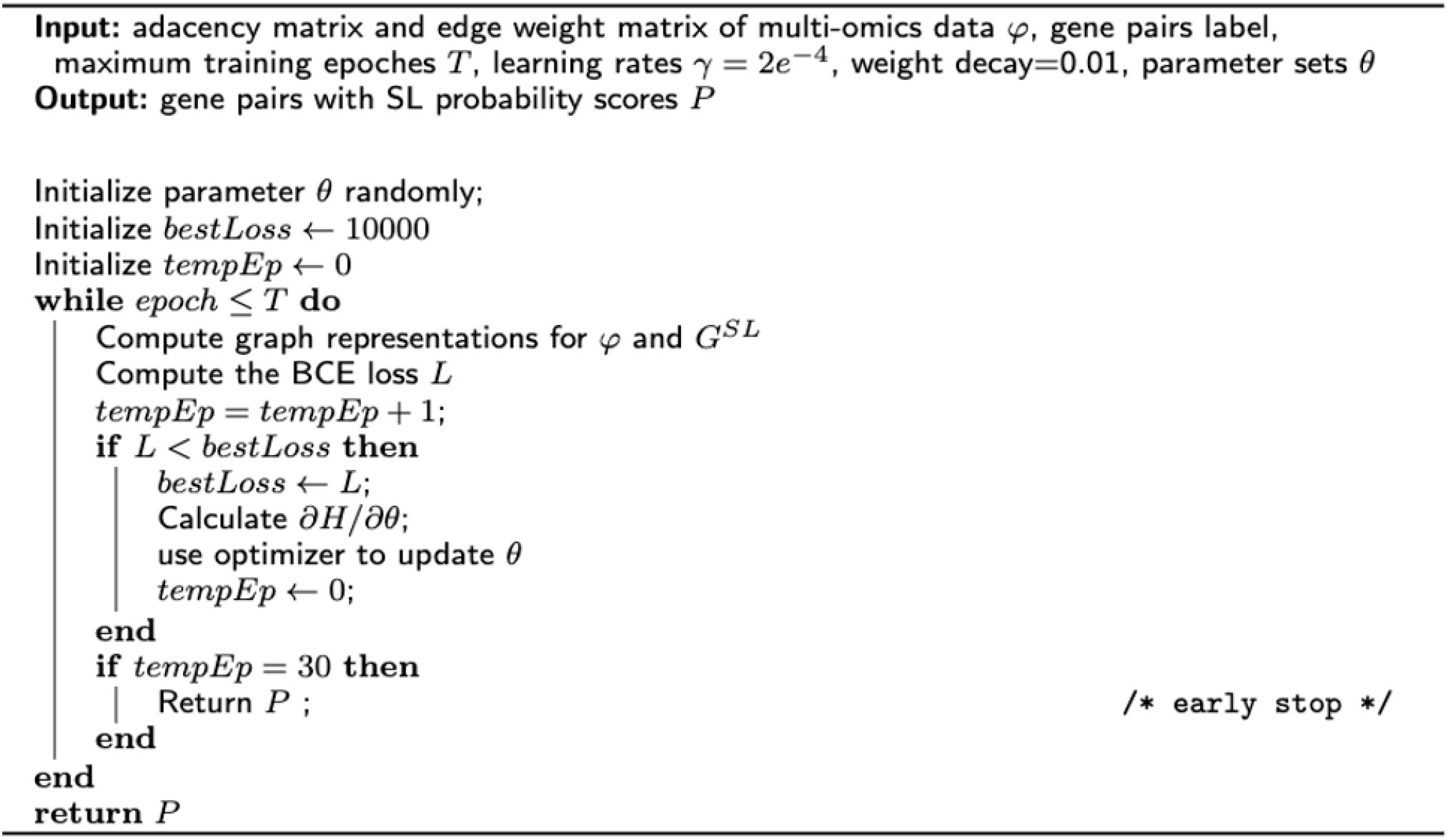

