## Supplemental Table1_SigProfiler_30-denovo-sigs for "Using graph-based model to identify cell specific synthetic lethal effects"

Table S1 The performance of architecture ablation test.

| Train Data | Test Data | Encoder | AUC | AUPR | Recall | Precision |
|---|---|---|---|---|---|---|
| A549 | A549 | SL_GCN | 0.888±0.03 | 0.887±0.05 | 0.867±0.30 | 0.404±0.37 |
| | | SL_GAT | 0.535±0.21 | 0.459±0.21 | 0.334±0.00 | 1.000±0.00 |
| | | SL_SAGE | 0.863±0.04 | 0.863±0.02 | 0.996±0.01 | 0.286±0.06 |
| | | SL_TAG | 0.604±0.09 | 0.586±0.05 | 0.836±0.37 | 0.164±0.11 |
| A375 | A375 | SL_GCN | 0.998±0.00 | 0.996±0.45 | 0.536±0.26 | 1.000±0.00 |
| | | SL_GAT | 0.520±0.10 | 0.449±0.21 | 0.335±0.00 | 1.000±0.00 |
| | | SL_SAGE | 1.000±0.00 | 1.000±0.00 | 1.000±0.00 | 0.922±0.07 |
| | | SL_TAG | 0.869±0.13 | 0.830±0.18 | 0.881±0.12 | 0.344±0.25 |
| HT29 | HT29 | SL_GCN | 0.870±0.09 | 0.893±0.08 | 0.432±0.19 | 0.947±0.12 |
| | | SL_GAT | 0.517±0.10 | 0.353±0.11 | 0.333±0.00 | 1.000±0.00 |
| | | SL_SAGE | 0.907±0.04 | 0.908±0.05 | 0.987±0.03 | 0.710±0.14 |
| | | SL_TAG | 0.654±0.08 | 0.519±0.10 | 0.692±0.20 | 0.169±0.08 |

Table S2 The performance of data ablation test

| Train Data | Test Data | Data combination | AUC | AUPR | Recall | Precision |
|---|---|---|---|---|---|---|
| A549 | A549 | EM+paralog+ES | 0.779±0.14 | 0.814±0.08 | 0.971±0.06 | 0.308±0.27 |
| | | EM+ES+L1000 | 0.840±0.12 | 0.850±0.09 | 0.985±0.03 | 0.311±0.26 |
| | | paralog+ES+L1000 | 0.805±0.12 | 0.817±0.10 | 0.971±0.06 | 0.312±0.26 |
| | | EM+paralog+L1000 | 0.838±0.12 | 0.849±0.09 | 0.983±0.04 | 0.299±0.23 |
| | | EM+paralog+L1000+ES | 0.863±0.04 | 0.863±0.02 | 0.996±0.01 | 0.286±0.06 |
| A375 | A375 | EM+paralog+ES | 0.933±0.01 | 0.937±0.01 | 1.000±0.00 | 0.664±0.07 |
| | | EM+ES+L1000 | 0.923±0.09 | 0.933±0.06 | 0.964±0.05 | 0.708±0.13 |

| | | | | | | |
|---|---|---|---|---|---|---|
| | | paralog+ES+L1000 | 0.923±0.06 | 0.944±0.05 | 1.000±0.00 | 0.738±0.13 |
| | | EM+paralog+L1000 | 0.923±0.11 | 0.947±0.08 | 1.000±0.00 | 0.723±0.21 |
| | | EM+paralog+L1000+ES | 1.000±0.00 | 1.000±0.00 | 1.000±0.00 | 0.922±0.07 |
| HT29 | HT29 | EM+paralog+ES | 0.835±0.01 | 0.714±0.02 | 0.712±0.10 | 0.594±0.10 |
| | | EM+ES+L1000 | 0.830±0.01 | 0.701±0.04 | 0.724±0.03 | 0.588±0.14 |
| | | paralog+ES+L1000 | 0.827±0.01 | 0.733±0.03 | 0.750±0.01 | 0.593±0.07 |
| | | EM+paralog+L1000 | 0.832±0.01 | 0.712±0.02 | 0.736±0.03 | 0.624±0.09 |
| | | EM+paralog+L1000+ES | 0.907±0.04 | 0.908±0.05 | 0.987±0.03 | 0.710±0.14 |

Table S3 The performance of attention module ablation test

| Train Data | Test Data | Split method | Method | AUC | AUPR | Recall | Precision |
|---|---|---|---|---|---|---|---|
| A549 | A549 | CV1 | attention | 0.863±0.04 | 0.863±0.02 | 0.996±0.01 | 0.286±0.06 |
| | | | No_attention | 0.788±0.09 | 0.771±0.12 | 0.915±0.09 | 0.446±0.32 |
| A375 | A375 | CV1 | attention | 1.000±0.00 | 1.000±0.00 | 1.000±0.00 | 0.922±0.07 |
| | | | No_attention | 0.833±0.04 | 0.782±0.12 | 0.766±0.05 | 0.692±0.13 |
| HT29 | HT29 | CV1 | attention | 0.907±0.04 | 0.908±0.05 | 0.987±0.03 | 0.710±0.14 |
| | | | No_attention | 0.815±0.18 | 0.855±0.14 | 0.959±0.09 | 0.694±0.24 |
